# conST: an interpretable multi-modal contrastive learning framework for spatial transcriptomics

**DOI:** 10.1101/2022.01.14.476408

**Authors:** Yongshuo Zong, Tingyang Yu, Xuesong Wang, Yixuan Wang, Zhihang Hu, Yu Li

## Abstract

**Motivation:** Spatially resolved transcriptomics (SRT) shows its impressive power in yielding biological insights into neuroscience, disease study, and even plant biology. However, current methods do not sufficiently explore the expressiveness of the multi-modal SRT data, leaving a large room for improvement of performance. Moreover, the current deep learning based methods lack interpretability due to the “black box” nature, impeding its further applications in the areas that require explanation.

**Results:** We propose conST, a powerful and flexible SRT data analysis framework utilizing contrastive learning techniques. conST can learn low-dimensional embeddings by effectively integrating multi-modal SRT data, *i*.*e*. gene expression, spatial information, and morphology (if applicable). The learned embeddings can be then used for various downstream tasks, including clustering, trajectory and pseudotime inference, cell-to-cell interaction, *etc*. Extensive experiments in various datasets have been conducted to demonstrate the effectiveness and robustness of the proposed conST, achieving up to 10% improvement in clustering ARI in the commonly used benchmark dataset. We also show that the learned embedding can be used in complicated scenarios, such as predicting cancer progression by analyzing the tumour microenvironment and cell-to-cell interaction (CCI) of breast cancer. Our framework is interpretable in that it is able to find the correlated spots that support the clustering, which matches the CCI interaction pairs as well, providing more confidence to clinicians when making clinical decisions.

## 1 Introduction

Single-cell RNA sequencing (scRNA-seq) technologies have enabled the characterization of the transcriptome of individual cells, yielding cell sub-populations across organs by high-throughput profiling. However, the dissociation step erases the spatial context of cells from their original tissue, which is of vital importance for understanding cellular functions and organizations.

Spatially resolved transcriptomics (SRT) (1) addresses these limitations by measuring the gene expression matrix and its corresponding spatial information simultaneously. Current spatial transcriptomics technologies can be mainly divided into two categories: (1) High-plex RNA imaging (HPRI) methods use probes targeting specific genes to localize mRNA transcripts. This method includes fluorescence *in situ* hybridization (FISH) or *in situ* sequencing (ISS), such as MERFISH (2), seqFISH (3), seqFISH+ (4), and STARmap (5). (2) Spatial labeling methods utilize spatial barcodes to capture mRNA transcripts across tissue cross-sections, and then deep sequencing is performed after detachment. Typical methods include ST (6), slide-seq (7), Slide-seqV2 (8), and 10x Genomics. The data generated from spatial labeling platforms, such as ST, 10x Genomics, are often accompanied by histology images (morphology) besides gene expression and spatial information. These two platforms are usually complementary: HPRI methods can achieve single-cell resolution with greater depth, while spatial labeling methods have greater coverage and are more accessible (9). In both platforms, mRNA transcripts are captured by the individual coordinates or spatial barcoding locations. For simplicity, we refer them to *spots* in the following texts.

The rich information from different modalities of the SRT data sheds light on the biological insights. A multitude of works utilizes Markov random field to integrate gene expression and spatial information. (10) proposed a hidden Markov random field (HMRF) model for cell type and spatial domain identification according to spatial dependency. Giotto (11) analyzes the correlation of gene expression among its neighbors for spatial domain detection. BayesSpace (12) utilizes the information of spatial neighborhood to iteratively update the model in a Bayesian manner. While these methods can achieve state-of-the-art performance on some specific tasks, such as cell type and spatial domain identification, they are unable to learn a universal embedding for downstream tasks, thus being less flexible.

Recently, there are some analytical tools that can generate low-dimensional embeddings. Seurat (13) adopts the single-cell analysis pipeline that is mainly focused on gene expression data while overlooking the spatial link. SpaCell (14) uses a ResNet (15) pretrained on ImageNet (16) to extract features from images of each spot and uses autoencoders to learn embedding from the extracted features and gene expression, where the spatial information is completely ignored. And as it is only tested on old version SRT data, it is unclear whether this method is effective for SRT of a higher resolution. stLearn (17) develops a Spatial Morphological gene Expression (SME) normalization method to recompute gene expression values by averaging neighboring spots. It also uses an ImageNet-pretrained ResNet to extract morphology features to calculate weights for averaging. The normalized expression values are used for downstream tasks, but containing excessive noise. SpaGCN (18) applies a graph convolutional network (GCN) to a graph constructed by spatial coordinates and morphological features, where gene expression of each spot is regarded as a node attribute. SpaGCN only incorporates the simple statistics of morphology during graph construction, and therefore the morphology may not be fully explored. SEDR (19) learns embeddings by reconstructing gene expression and spatial information with an autoencoder and a variational graph autoencoder, respectively. Morphological information is ignored in SEDR pipeline.

To sum up, there are three major limitations that remain unsolved in the previous methods. (1) The morphological features are extracted by pretrained CNN or simply ignored. It has not been involved in the training process for further integration. (2) The biological relationship between spots, sub-clusters, and the global structure of the SRT data has not been fully explored during the embedding learning process, remaining a large room for performance improvement. (3) While the learned embeddings can be used in various downstream tasks, the “black box” model stems the interpretation of the obtained results.

Here, we present an interpretable multi-modal contrastive learning framework for spatial transcriptomics, conST, to address the above-mentioned problems. For the first concern, conST can effectively integrate gene expression, spatial information, and morphology (if accessible) to learn low-dimensional embeddings. A state-of-the-art computer vision model, MAE (20), is used to extract informative features from the morphology of each spot. The extracted features are then regarded as an individual node attribute to participate in the training. Our framework is also very flexible. When morphology is not available, it can take gene expression and spatial information as input, thus can be applied for both HPRI and spatial labeling data.

As for the second concern, we argue that there are three natural underlying relationships in SRT data that can be used as supervision signals to guide the network to learn more meaningful embeddings: (1) local: a small portion of noises does not prevent a spot to be identified by its distinguishing features, (2) global: as a dataset is taken from the same slice (*e*.*g*., same piece of tissue), the spots within it possess similar globally general features, (3) context: the node features are more related inside a sub-cluster, *e*.*g*., more similar expression pattern and imaging texture. Based on the above assumptions, we propose to utilize contrastive learning (21; 22) to learn embeddings by maximizing the mutual information in local-local, local-global, and local-context levels.

In terms of interpretation, we monitor the training process and utilize GNNExplainer (23) to identify the important subgraphs and node attributes contributing most to the predictions according to the mutual information. Therefore, we can not only obtain the results, but also know what contributes to the prediction, *i*.*e*., finding the correlated spots that support the clustering, which matches the CCI interaction pairs as well. The interpretability will give clinicians more confidence when using conST for the real clinical data analysis. We demonstrate how to analyze the tumour microenvironment and cell-to-cell interaction of breast cancer tissue with conST, leading to a clear cancer progression prediction, which will be helpful for making clinical treatment decisions.

The experimental results demonstrate that the learned embeddings can be successfully applied to many downstream tasks, such as clustering, trajectory inference, spatially variable genes (SVGs) detection, batch correction, and cell-to-cell interaction, *etc*. Quantitatively, conST outperforms other concurrent methods, *e*.*g*., by up to 10% increase regarding ARI on clustering tasks. The qualitative results in both HPRI and spatial labeling platforms demonstrate the superiority of our method.

conST is user-friendly and flexible. With the standard SRT input, users can obtain the embeddings in the commonly-used formats in an end-to-end manner. We also provide detailed tutorials for downstream analysis with the generated embeddings.

Our main contribution can be summarized as follows:

- We propose conST, a powerful and flexible multi-modal framework that can effectively incorporate gene expression, spatial information, and morphology to learn low-dimensional yet expressive embeddings for downstream tasks.
- To the best of our knowledge, we are the first to introduce contrastive learning in the area of spatial transcriptomics analysis, demonstrating its effectiveness in exploring the pattern from the data itself.
- The use of GNNExplainer enhances the interpretability of conST, giving users more confidence especially when used in clinical situations by not only telling what, but also why.
- Extensive experiments are conducted using the learned embedding in various datasets. The experimental results demonstrate the superiority and robustness of the proposed method in both HPRI and spatial labeling platforms.

## 2 Methods

### 2.1 Problem Definition and Framework

The whole framework is presented in Figure 1. Gene expression is undoubtedly the deterministic factor of the biological property. Besides, the rich information contained in histology images has been proven to be successful in revealing other important characterizations (24; 25). The spatial information provided by SRT acts as a bridge linking them together so that gene expression and morphology can complement each other as well as be more aware of the neighborhood.

**Figure 1:**
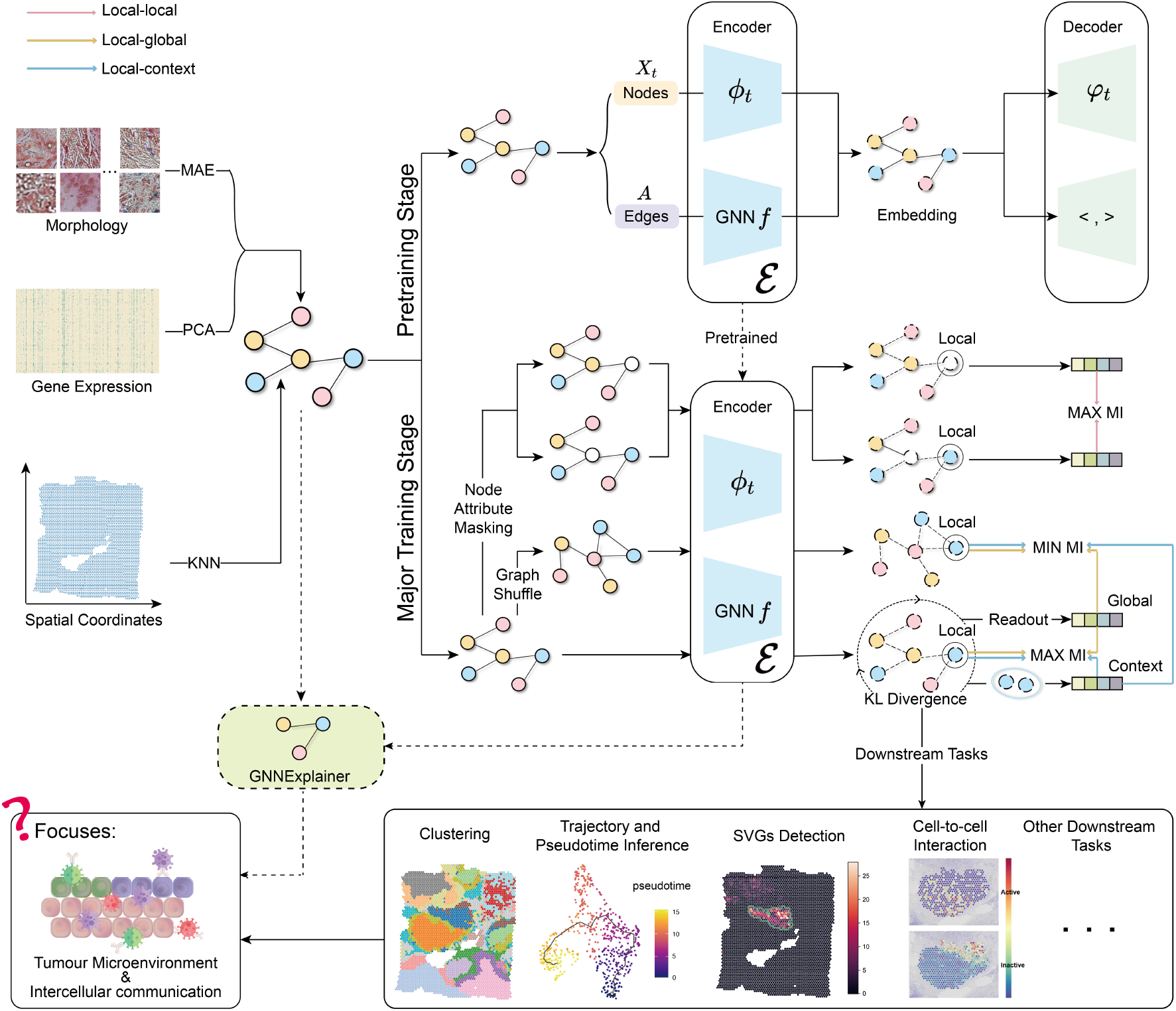
Framework of conST. conST models the ST data as a graph by treating gene expression and morphology as node attributes and constructing edges by spatial coordinates. The training is divided into two stages: pretraining and major training stage. Pretraining stage initializes the weights of the encoder *ℰ* by reconstruction loss. In major training stage, data augmentation is applied and then contrastive learning in three levels, *i*.*e*., local-local, local-global, local-context, are used to learn a low-dimensional embedding by minimize or maximize the mutual information (MI) between different embeddings. The learned embedding can be used for various downstream tasks, which, when analyzed together, can shed light on the widely concerned tumour microenvironment and cell-to-cell interaction. GNNExplainer helps to provide more convincing predictions with interpretability.

Therefore, naturally, their underlying relationship can be modeled by graph structure and processed by graph neural networks (GNNs) (26). Hence, we treat the embedding generation process as a multi-modal self-supervised graph representation learning problem. In our setting, a graph is constructed based on K-Nearest Neighbor (KNN) distance of the coordinates of each spot, where the spots are regarded as nodes whose attributes are gene expression and morphology features (if accessible).

Denote 𝒢 = (𝒱, *E*) as a graph with *N* nodes, where 𝒱 = {*v*_1_, *v*_2_, …, *v*_*N*_ } represents the node set and *E* ⊆ 𝒱 × 𝒱 represents the edge set. The feature matrix of all the nodes is denoted as **X** = {*x*_1_, *x*_2_, …, *x*_*N*_ } ⊆ ℝ^*N×F*^, where *F* is the dimension of node features. If nodes possess multiple attributes, the *t*-th feature matrix is denoted as 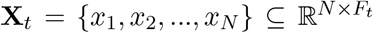. The adjacency matrix is given by **A** = [0, 1]^*N×N*^, of which **A**_*ij*_ = 1 if {*v*_*i*_, *v*_*j*_} ⊆ *ℰ* or 0 otherwise.

Our objective is to learn a general encoder *ℰ*(**X, A**) in a self-supervised manner that can produce node embeddings in the low-dimensional space. Denote the learned embedding **H** = *ℰ*(**X, A**) ⊆ ℝ^*N×F ′*^, where *F*^*′*^ ≪ *F*, and **h**_**i**_ is the embedding of node *v*_*i*_. We utilize Graph Convolutional Networks (GCNs), a powerful variants of GNN. Let **Â** denote the normalized adjacency matrix and **H**^(*l−*1)^ denote the embedding of layer *l* − 1. The propagation of the GCNs is defined as **H**^(*l*)^ = *σ*(**ÂH**^(*l−*1)^**W**^*l*^), where **W**^*l*^ is a learnable weight matrix and *σ* is a non-linear activation function. The normalized adjacency matrix is defined as 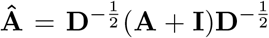, where **D** is the degree matrix of **A**, *i*.*e*., **D**_*ii*_ = Σ_*j*_ **A**_*ij*_, and **I** is an identity matrix for self-loop.

The training can be divided into two stages. In the pretraining stage, gene expression and morphology features are input into an autoencoder-based network to further reduce dimensions and initialize the weights of the encoder *ℰ*. In the major training stage, we focus on contrastive learning to guide the network to learn more informative and robust embeddings.

### 2.2 Pretraining stage

Based on the euclidean distance, the adjacency matrix **A** is constructed, where **A**_*ij*_ = 1 if {*v*_*j*_} is in the KNN of {*v*_*i*_} and 0 else. We filter the genes that have very low expression and use principle component analysis (PCA) to reduce the expression matrix to dimention *F*_1_, *i*.*e*., 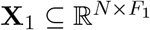.

The morphological features of each spots are extracted by a pretrained MAE model (20), which achieves SOTA performance and transferability on various datasets. Due to the small number of the available Hematoxylin and Eosin (H&E) stain images from ST, MAE is pretrained using ImageNet in a self-supervised manner. The recent development in computer vision communities has proven that features extracted by self-supervised models are more robust and transferable (27). MAE contains an asymmetric encoder-decoder architecture, with a Vision Transformer (ViT) (28) encoder. The decoder is used for pre-training, and only its encoder is used to extract the morphology features. The pre-training process masked a large proportion (75%) of each image for reconstruction with positional encoding to enable the model to be aware of both the global and local information, while the large ViT makes the extractor more powerful. The detailed explanation of MAE can be found in the supplementary material. Compared with supervised pretrained ResNet used in previous methods (14; 17), we argue that the MAE is more powerful for extracting features of the H&E stain images. The extracted feature dimension of each spot is *F*_2_, and the feature matrix is regarded as another node attribute 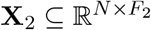.

In the pretraining stage, we take advantage of proximity-based learning to initialize the parameters and further reduce the dimension of node attributes. For node attribute 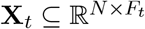, a deep autoencoder Φ_*t*_ with an encoder *ϕ*_*t*_ and decoder of *φ*_*t*_ is used. The encoder produce a latent embedding 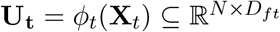, where *D*_*ft*_ denotes the dimension of the learned low-dimensional embedding. After that, a variational graph autoencoder (VGAE) (29) with a encoder *f* is used to encode the spatial information into the node features. Depending on whether morphology is available, denote **U** = ∥_*t*_**U**_*t*_, where ∥ is the concatenation operation. The latent embedding from the VGAE is obtained by 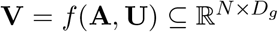. The final embedding **H** is the concatenation of **U** and **V** ⊆ ℝ^*N×F ′*^, where *F*^*′*^ = Σ_*t*_ *D*_*ft*_ + *D*_*g*_.

To be more specific, the encoder *f* of the VGAE is a two-layer GCN whose inference process is defined as

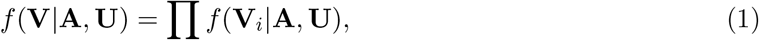

with

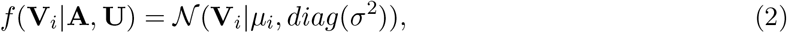

where the ***μ*** = GCN_*μ*_(**A, U**) is the matrix of mean vectors ***μ***_*i*_, log *σ* = GCN_*σ*_(**A, U**), and 𝒩 is the normal distribution.

The decoding part of the autoencoder Φ_*t*_ and VGAE all take the concatenated embedding **H** as input, which possesses both spatial information and node attributes. For the autoencoder Φ_*t*_, the decoder *φ*_*t*_ tries to reconstruct the input feature matrix as 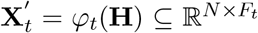. The decoder of VGAE is given by a simple inner product to generate the reconstructed **A**^*′*^ = *σ*(**H · H**^*T*^).

The autoencoder Φ_*t*_ is optimized by Mean Squared Error (MSE) between **X**_*t*_ and 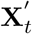. The VGAE is optimized by minimizing the standard cross entropy between the input **A** and reconstructed **A**^*′*^, as well as Kullback-Leibler (KL) divergence between *f* (**H**|**A, U**) and the Gaussian prior *p*(**H**) = Π_*i*_ *p*(**h**_*i*_) = Π_*i*_ *𝒩* (**h**_*i*_|0, **I**).

### 2.3 Major training stage

In the major training stage, the encoder part of the autoencoder and the VGAE are preserved, and the decoder part is dropped. We hope to learn a general encoder *ℰ* = {*ϕ*_*t*_, *f*} that produce the embeddings, *i*.*e*., **H** = *ℰ* (**X, A**). Contrastive learning is then used to supervise the network for better representation learning at three levels: local-local, local-context, and local-global.

#### Local-local

In the process of sequencing, noises to the gene reads are commonly introduced by device deficiencies or manual operations. However, experts are still able to identify important characteristics of a specific spot based on the overview of its gene expression and the marker genes. Similarly, we hope the network to be able to distinguish a specific node from other nodes even if there are unwanted noises. Motivated by this and the recent work in node-level graph contrastive learning (30), the local-local level contrastive learning is utilized to force the network to focus on more important features of the node attributes. Two views of the original graphs are created by masking node attributes. Then, the mutual information of the same node between these two views is maximized.

Specifically, we first randomly mask the node attributes of each spot at a given ratio to construct two augmented graphs 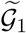 and 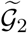, with an unchanged adjacency matrix **A** and masked feature matrices 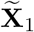 and 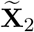. The masked feature matrix 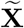 is computed by

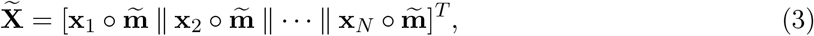

where ∥ is concatenation operation and ° is the element-wise product. 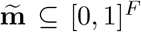 is a random sampling vector. Given a node dropping probability *p*_*m*_, each dimension of 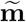 is independently drawn from a Bernoulli distribution, namely, 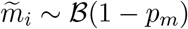.

Denote generated embeddings from the two augmented graph as **H**_**U**_ and **H**_**V**_. The contrastive objective is to distinguish the embedding of the same node in two augmented views from that of the other nodes. For node *v*_*i*_, we set its embedding 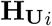 generated in one view as an anchor. The embedding 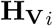 generated in the other view is treated as a positive sample, and the rest of nodes are treated as negative samples. Define a similarity measurement as sim(**H**_**U**_, **H**_**V**_) = *θ*(*p*(**H**_**U**_), *p*(**H**_**V**_)), where *θ* is the cosine similarity and *p*(·) is a projection head constructed by a two-layer multiple layer perceptron (MLP). Formally, for a positive pair 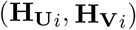 of node *v*_*i*_, the objective is formulized similar to NT-Xent (31). For positive pair, we have 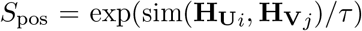, where *τ* is the a temperature hyperparameter. For negative pairs within a same view, *i*.*e*., intra-view, we have 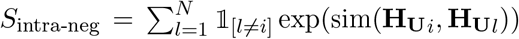, where 𝕝_[*l ≠ i*]_ ⊆ {0, 1} is the indicator function. And for negative pairs between two views, *i*.*e*., inter-view, we have 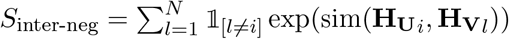. Thus, the objective can be given by

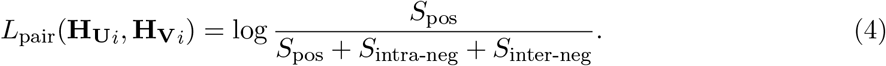

Since the two views are symmetric, the whole objective for local-local contrastive learning is calculated by averaging all positive pairs as

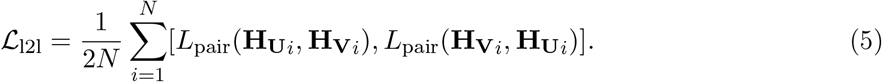

#### Local-global

For a specific SRT dataset (*e*.*g*., a slice), the gene expression or morphology of spots are intrinsically coherent with the global property, as they are captured from the same section of the same species. Therefore, it is beneficial to maximize the mutual information between the embeddings of each individual node and the whole graph summary to endow learned embeddings with the global structure and more robustness to the neighboring noises.

We adopt a similar method with Deep Graph Infomax (DGI) (32), a commonly used graph self-supervised method that has shown superior performance. First, a *readout function* ℝ is used to obtain the overall summary **s** of the graph, where ℛ : ℝ^*N×F ′*^ → ℝ^*F ′*^ and **s** = ℛ (ℰ(**X, A**)). A corrupted graph 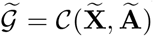 is obtained by a corruption function𝒞, which randomly shuffles the rows of the feature matrix and randomly adds/drops edges. The embeddings obtained from original graph and corrupted graph are denoted as **H** and 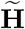.

A discriminator 𝒟_*G*_ : ℝ^*F ′*^ × ℝ^*F ′*^ is employed to assign higher probability scores to positive pairs (**s, h**_*i*_) than the negative pairs 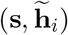. The objective is to maximize the Jensen-Shannon-based BCE as

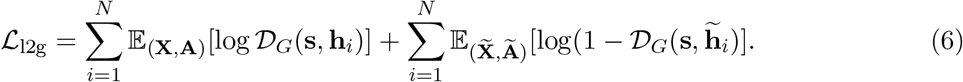

#### Local-context

Besides the global property, spots tend to be more similar at the cluster level. For example, spots may share similar marker genes and morphological texture within the same tissue type. Hence, we also try to maximize the mutual information between the node attributes and the cluster-level summary, which is beneficial for learning the embedding with hierarchical structure.

Inspired by Graph InfoClust (33), first, we use a differentiable K-Means clustering algorithm (34) to obtain *M* clusters. The centroid of each clusters ***μ***_*m*_ ⊆ ℝ^1*×F ′*^ is updated iteratively through a cluster layer

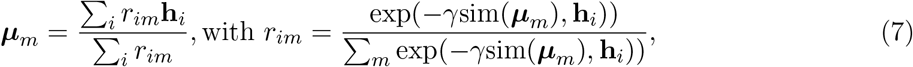

where *m* = 1, 2, …, *M*, and *γ* is a inverse-temperature hyperparameter. For each node, we maximize the mutual information between **h**_*i*_ and its corresponding cluster summary **z**_*i*_. **z**_*i*_ is computed as

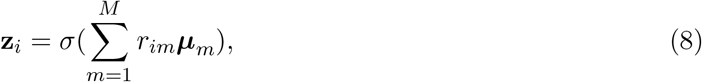

where *r*_*im*_ is the probability of node *v*_*i*_ assigned to cluster *m* and 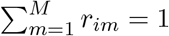.

Similar to local-global level, another discrimintor 𝒟_*C*_ is used to assign higher scores to the positive pairs (**h**_*i*_, **z**_*i*_) than the negative pairs 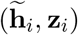. Note that we use the same corruption function 𝒞 as that in the local-global level. The objective is also to maximize the Jensen-Shannon-based BCE as:

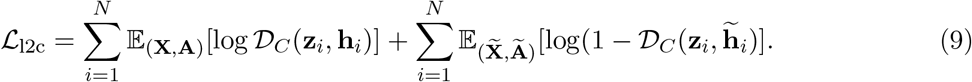

Weighted by *λ*_*i*_, the overall contrastive learning objective function is given by

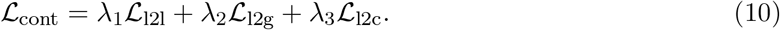

#### KL Divergence

In addition to contrastive learning, we use a deep clustering method (35) to further refine the embedding for more compactness. Simply, we use the normal K-means to cluster the embedding to *K* clusters. First, a Student’s t-distribution kernel is used to calculate the soft assignment probability *q*_*ij*_ of the embedding **h**_*i*_ to the cluster centroid ***ν***_*j*_

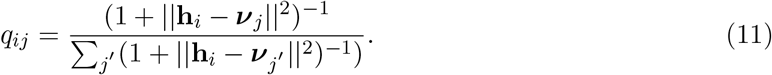

Next, based on *q*_*ij*_, a target distribution *P* is calculated to help learn from the assignments with higher scores

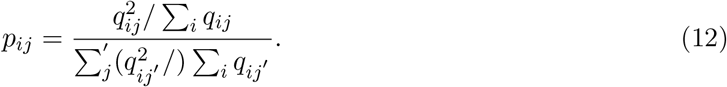

Finally, an auxiliary KL Divergence objective is defined as

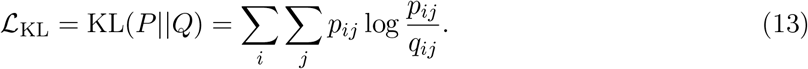

Finally, the loss function in the major training stage is

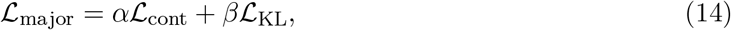

where *α* and *β* are weight factors.

### 2.4 GNNExplainer

GNNExplainer (23) is a method for providing explanations for any GNN-based models in a model-agnostic manner. Here, we use the trained conST model and specific node index we would like to inspect as inputs. The GNNExplainer can identify an important subgraph and node features that are most influential to the predictions as an explanation by maximizing the mutual information between the subgraph and the predictions.

Formally, for a given node *v*_*i*_, GNNExplainer aims to identify a subgraph 𝒢_*S*_ ⊆ 𝒢 and the corresponding feature matrix **X**_*S*_ = {**x**_*j*_ |*v*_*j*_ ⊆ 𝒢_*S*_ }. The GNNExplainer can be optimized according to the mutual information (MI) as:

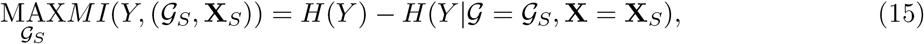

where *Y* is the prediction by conST, and *H*(), *H*(|) are the marginal entropy and conditional entropy, respectively. The mutual information measures whether removing an edge link or a node feature is determinant to the final prediction results. Thus, by maximizing the MI, the GNNExplainer returns a subgraph and node features that have the highest mutual information with the original input, *i*.*e*., the returned subgraph and node features can lead to the most similar prediction as the original input.

For conST, we are not only interested in what predictions it makes, but also why it makes such predictions. The GNNExplainer can identify which spots have the most important correlation with the given spot and their correlation can further explain gene expression and cell-to-cell interaction activities.

## 3 Experiments

### 3.1 Datasets and Implementation Details

We use various datasets from both HPRI and spatial labeling platforms to verify the effectiveness of our method. For HPRI data, we use mouse hypothalamus MERFISH (36), mouse visual cortex seqFISH (10). For spatial labeling data, we use human dorsolateral prefrontal cortex spatialLIBD (37) generated by 10x Visium, Human Breast Cancer (Block A Section 1) generated by 10x genomics (38), and mouse olfactory bulb Stereo-seq (39).

We set *K* = 20 for KNN graph construction. PCA is used to reduce the dimension of gene expression to *F*_1_ = 300. The dimension of morphological features is *F*_2_ = 768. Users can also easily adjust these parameters accordingly to meet the requirements of specific datasets. The two stages are trained 200 epochs respectively.

For downstream tasks, Leiden algorithm (40) is used for clustering; Monocle3 (41) and PAGA (42) is utilized for trajectory inference and pseudo-time inference; we adopt a similar SVGs detection method as SpaGCN (18) with our own embedding; Cell-to-cell interaction is performed with TraSig and Seurat (13) for label transfer. The detailed descriptions of datasets and implementation can be found in the supplementary material.

### 3.2 Clustering

Accurate clustering of spatial domains or cell types reveals the structure of the SRT data. We experimented on SRT data produced in different platforms with different resolutions, coverages, and depths. The results prove that conST is a powerful and generalizable framework that can be used to cluster both HPRI and spatial labeling data.

As shown in Figure 2, we compare the clustering results of conST with other methods in spatialLIBD dataset and illustrate slice 151673 which has clear layer boundaries. Visually, the results of conST do not have distinct outliers compared to other methods. Also, although the results of SpaGCN, SEDR, and BayesSpace all have seven layers that are relatively clear, the white matter (WM) layer and Layer 1 together (the first two layers in left bottom of the ground truth) they clustered are of the similar size of the WM layer in the ground truth. That will lead to the mismatch of the following layers, while each of the cluster boundaries of conST aligns well with the ground truth. conST achieves an ARI of 65%, about 10% higher than the state-of-the-art method. Also, conST achieves the highest mean and median ARI among all the methods.

**Figure 2:**
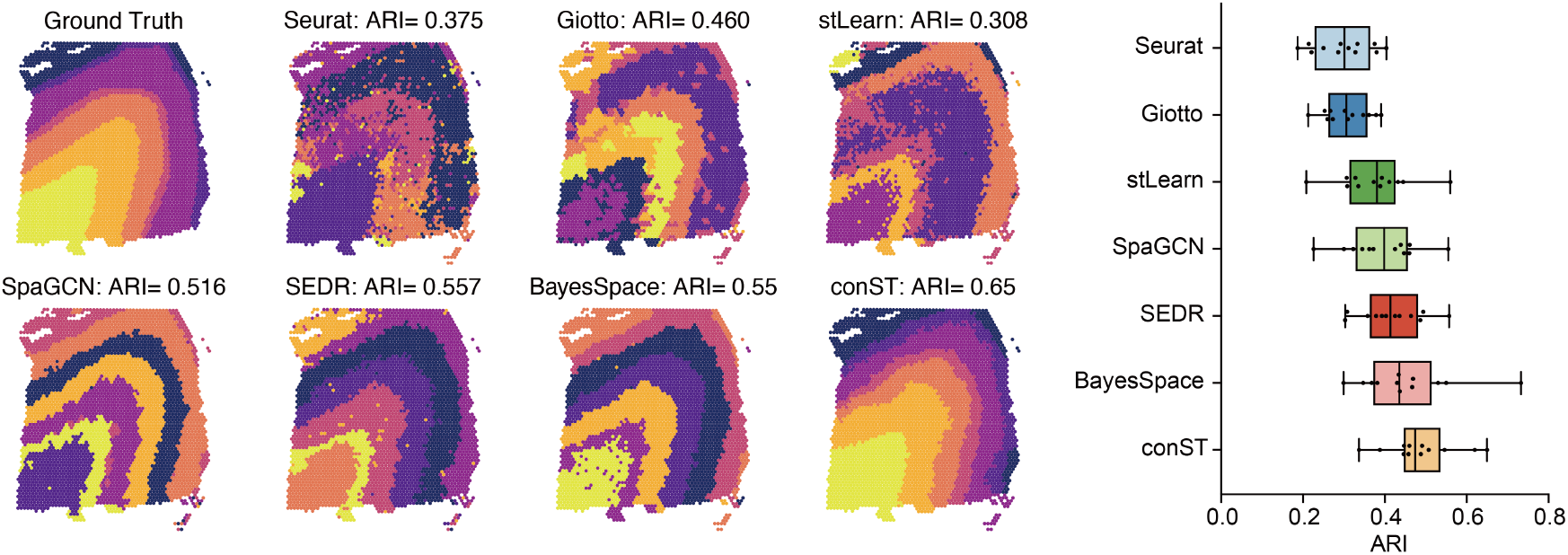
Comparison of different methods on slice 151673 of spatialLIBD dataset evaluated by ARI. conST achieves an ARI of 0.65, around 10% increase compared to other state-of-the-art methods. conST also achieves the highest mean and median ARI over all the 12 slices.

For HPRI data, we experimented with MERFISH and seqFISH data. Different from spatial labeling data, they are captured in a higher resolution but with fewer genes in each spot and no morphology is available. Even so, conST still demonstrates strong performance. In Figure **??**, we compare the clustering performance of mouse hypothalamus MERFISH data with SpaGCN and SEDR. Visually, conST produces more consistent clusters. Quantitatively, as there is no ground truth for MERFISH data, we use three unsupervised metrics to evaluate the quality of the clusterings, *i*.*e*., Silhouette Coefficient (SC), Calinski Harabasz Score (CHS), and Davies Bouldin Index (DBI) (Supplementary material Eq S1, S2, S6). For SC and CHS, the higher the better, while for DBI, the lower the better. conST significantly outperforms SpaGCN and SEDR on all the three metrics.

We also evaluated conST on Stereo-seq and seqFISH data (Supplementary Figure S2, S3), where conST produces clear boundaries and biologically meaningful clusters. The statistical results also prove the superiority of our method.

#### Ablation Study

Clustering is arguably the most intuitive way to justify the quality of the learned embeddings. We perform an ablation study to verify the necessity and effectiveness of each of the contrastive learning components. Results on the spatialLIBD dataset demonstrate that all of the three levels of contrastive learning are beneficial to embedding learning. It can be seen from Table 1 that the local-local level has the most important influence on the performance, of which the data augmentation randomly masks some features of the node attributes. It produces a similar effect as dropout (44) that widely occurs in SRT data as well. Then, by using contrastive learning, conST can effectively learn to distinguish the more important invariant features among the noising data, thus obtaining a stronger ability. It achieves the best performance when all of the three contrastive losses are used together, as shown in Table 1.

**Table 1:** Ablation study of different contrastive learning components measured by ARI.

| Case | Slice 151673 | Mean | Median |
| --- | --- | --- | --- |
| w/o local-local | 0.559 | 0.410 | 0.421 |
| w/o local-global | 0.615 | 0.457 | 0.438 |
| w/o local-context | 0.586 | 0.423 | 0.419 |
| All | **0.650** | **0.487** | **0.474** |

### 3.3 Trajectory and Pseudotime Inference

Trajectory inference infers the pattern of cells in the dynamically developmental progress, and pseudotime represents the progression through this progress. Monocle3 (41) and PAGA (42) are used to produce trajectory and pseudotime inference where we replace the default PCs with the generated embeddings to validate the quality of the learned embeddings. For pseudotime in the spatialLIBD dataset, white matter (WM) is selected as the starting point.

As shown in Figure 4a, the trajectory generated from our embeddings are more consistent along with the clusters, covering almost all the spots, while the embeddings of SpaGCN are rewinded like a circle and the trajectory of SEDR does not reach Layer 6. Also, our pseudotime is smoother with much fewer outliers than the other two methods.

**Figure 3:**
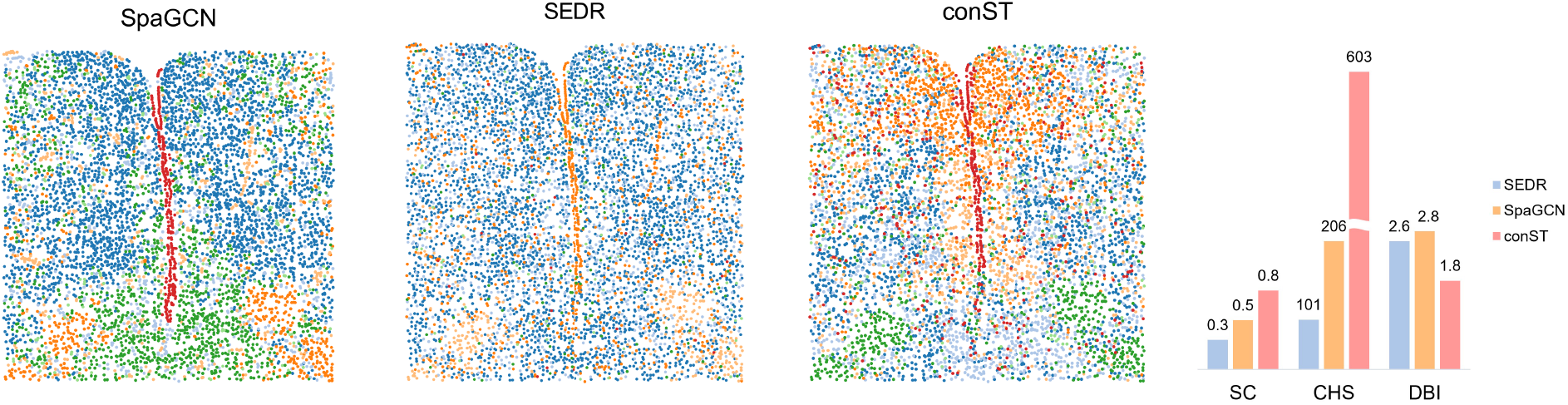
Comparison of the clustering results on MERFISH data. conST produces visually more consistent clusters and outperforms SpaGCN and SEDR in Silhouette Coefficient (SC), Calinski Harabasz Score (CHS), and DaviesBouldin Index (DBI). SC and CHS are the higher the better, while DBI is the lower the better.

**Figure 4:**
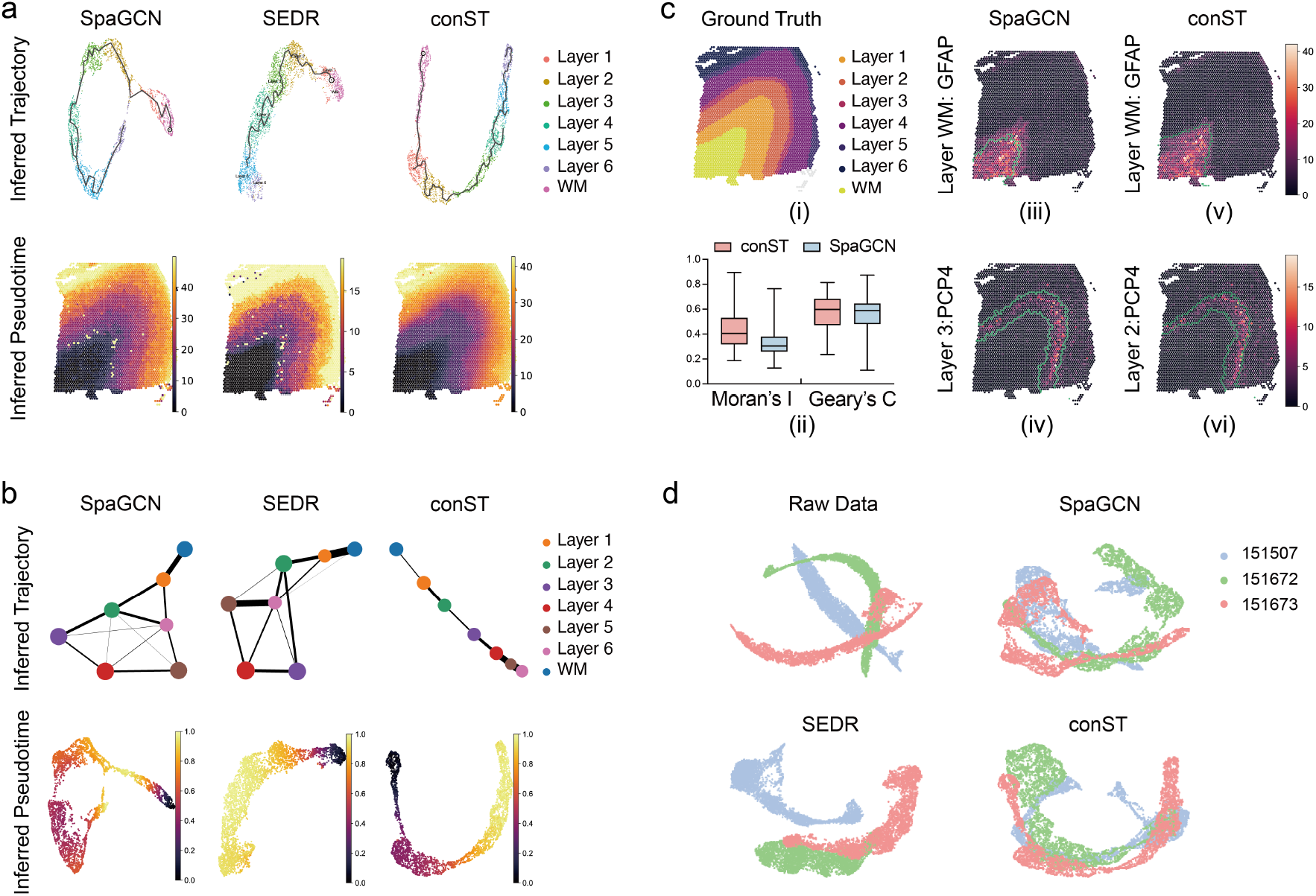
**a**. Comparison of trajectory and pseudotime inference using Monocle3 based on the learned embeddings of SpaGCN, SEDR, and conST. The trajectory inferred from conST goes through all the layers consistently and the pseudotime is much smoother with fewer outliers. **b**. Comparison of PAGA trajectory and pseudotime inference based on the learned embeddings of SpaGCN, SEDR, and conST. The trajectory inferred from conST is nearly linear, showing a clear progression process. The pseudotime of conST is also more consistent compared to SpaGCN and SEDR. **c**. SVGs detection comparison of SpaGCN and conST. conST outperforms SpaGCN in both Moran’s I and Geary’s C. The expression pattern of SVGs detected by conST aligns better with the actual anatomical layers, where the green lines indicate the predicted clustering boundaries. **d**. conST can alleviate batch effects. Raw data have apparent batch effects. Compared with other methods, conST can correct batch effects while preserving the semantic meanings.

In Figure 4b, the trajectory of conST inferred from PAGA is nearly linear, indicating a clear progression process from the white matter to the layer 6. It is also worth noticing that there are connections between layer 4 and layer 6 in the trajectory infered from conST, where the layer 4 to layer 6 are thin and close to each other in the ground truth. It is possible that they are not well diverged, indicating the biological consistency of our prediction. The trajectories from SpaGCN and SEDR are disordered with crossing lines between different layers. We also visualized the pseudotime on embeddings projected by UMAP. conST shows more consistent progression than SpaGCN and SEDR.

Furthermore, the accurate trajectory and pseudotime inference by conST can be also beneficial for more in-depth analysis such as cell-to-cell interaction, as detailed in Section 3.6.

### 3.4 SVGs Detection

Spatially variable genes (SVGs) have different expression patterns in different spatial locations. Detection of SVGs is helpful for identifying the tissue structure and clinical phenotypes. The better clustering performance enables conST to detect SVGs more accurately. For a fair comparison, we use the same detection method and default parameters as that in SpaGCN.

For slice 151673 in spatialLIBD, we have better performance on two commonly used metrics Moran’s I and Geary’s C (Figure 4c (ii)), which demonstrates SVGs detected by conST have a higher spatial autocorrelation. Furthermore, we select gene GFAP and PCP4 for comparison, as they are all highly expressed and detected by both SpaGCN and conST. While GFAP is detected in Layer WM by both methods, it can be seen that the boundary of Layer WM of conST aligns well with the highly expressed area of GFAP than SpaGCN (Figure 4c (iii), 4c (iv)), compared with the ground truth (Figure 4c (i)). For PCP4 (Figure 4c (v), 4c (vi)), although SpaGCN and conST all have cluster boundaries aligned well with the highly expressed area, it is however detected at Layer 3 by SpaGCN, which should be in Layer 2 as shown in ground truth and conST. The wrong layer could even lead to worse understanding if we want to know the relationship between the layers and SVGs. Hence, conST can be a more accurate tool for SVGs detection, and it confirms the superiority of our clustering results in turn.

### 3.5 Batch Correction

The batch effect refers to the changes to the result data in different experiments caused by non-biological factors, which can lead to imprecise conclusions. Batch effects are commonly observed in high-throughput sequencing experiments including SRT data. conST is able to alleviate the batch effects through contrastive learning. The local-global level contrastive learning enables spots to be aware of the global property. In this way, the learned embeddings of different slices of the same species will share similar properties. Furthermore, the local-local level contrastive learning makes the network more robust to the technical noises introduced by devices or manual operations (45; 46). Thus, the learned embedding can be projected into a shared latent space, correcting batch effects. We select three slices showing substantial batch effect (19) in spatialLIBD dataset for illustration.

We project the learned embeddings to 2-D space by UMAP (47). As shown in Figure 4d, the raw data demonstrate distinct batch effects. conST effectively corrects the batch effects with more compact and mixed embeddings, compared with SpaGCN and SEDR.

### 3.6 Cell-to-cell Interaction

For a tumour tissue, normally it can be grouped into 4 main morphotypes: Ductal Carcinoma in Situ (DCIS), healthy tissue (Healthy), Invasive Ductal Carcinoma (IDC), and tumor surrounding regions with low features of malignancy (Tumor edge) (48). DCIS consists of the proliferation of malignant cells which do not invade the basement membrane of the breast ducts. It is a nonobligate precursor to IDC and a large amount of DCIS lesions remain indolent in practice. However, currently, almost all the DCIS lesions are treated in practice to prevent further invasion, and the treatment comprises either mastectomy or breast-conserving radiotherapy surgery that will be harmful to the patients (49).

To counter the overtreatment, we predict the potential cancer progression by conST, hoping to provide more insights for clinical decision-making. First, the clinically potential target receptors on breast cancer cells are detected. Then, we analyze the neighbor-spreading trend for the IDC region and evaluate the risk for developing invasive breast cancer with a tool of cell-to-cell interaction (CCI).

We obtained the embeddings of the dataset of Human Breast Cancer (Block A Section I) (38) with conST, in short as BRCA. With the learned embeddings, we cluster them into 20 regions (Supplementary Figure S4). Given the heterogeneity of the tissue, we specifically focus on the obvious lesion area, *i*.*e*., cluster c2, c6, and c10 (Figure 5b (i)). Trajectory inference is applied to these three clusters to gain pseudotime ordering (Supplementary Figure S10) as we did in section 3.2 and 3.3. Then, by detecting SVGs, we find some highly expressed genes, including IGFBP2, PVALB, RABEP1, RAB11FIP1, LINC00645, *etc*. (Supplementary Figure S8, S9).

**Figure 5:**
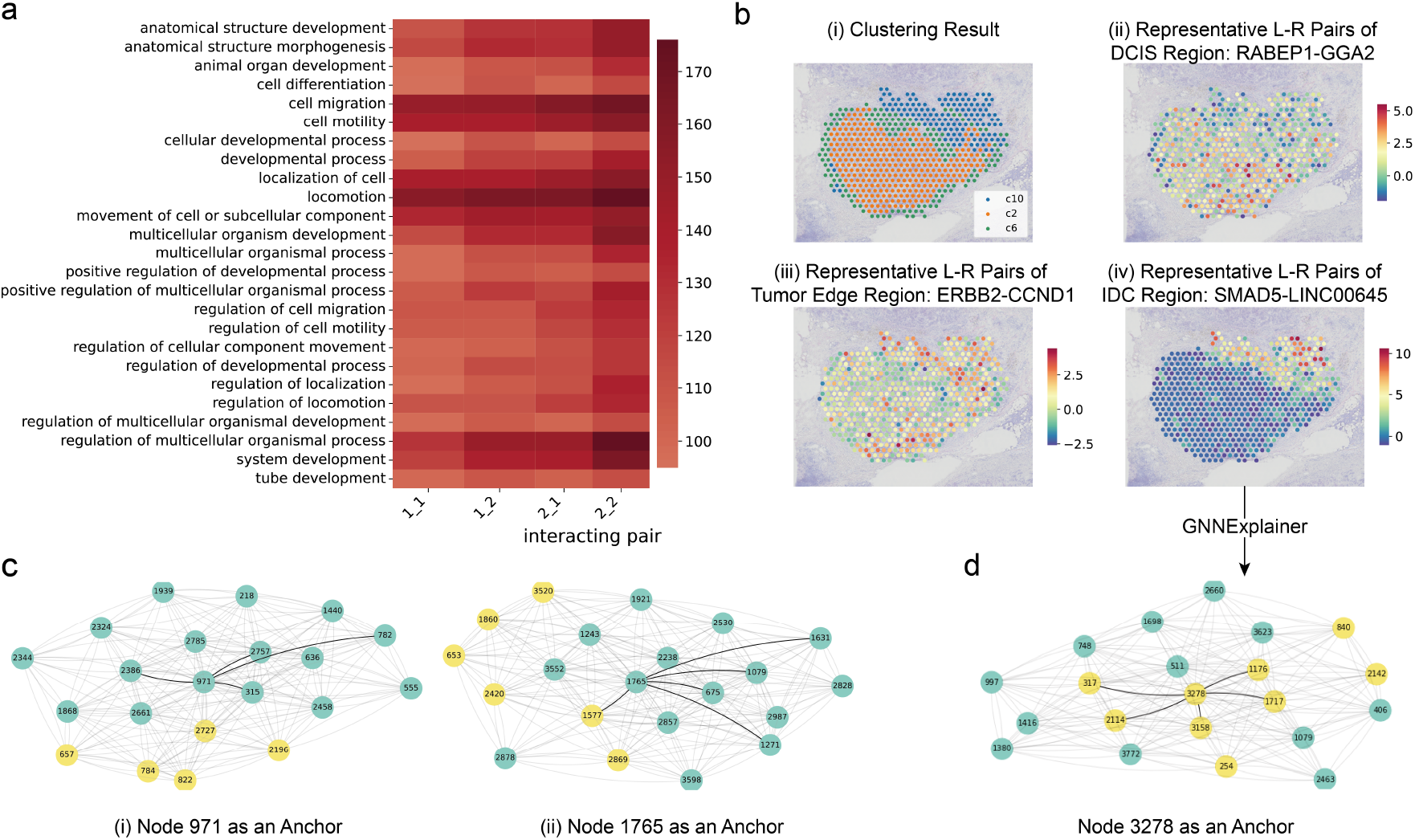
Cell-to-cell interaction and GNNExplainer. **a**. GO term analysis for cluster c2, c6 and c10 using Trasig based on embedding from conST. **b**. Representative L-R pairs of cell-to-cell interaction analysis. (i) Three clusters obtained by conST. (ii) (iii) (iv) representative L-R pairs of DCIS, Tumor edge, and IDC region respectively. **c**. GNNExplainer can identify subgraphs that contribute most to the predictions. Node 971 and 1765 in cluster boundary are selected as anchor nodes for illustration, whose identified subgraphs are plotted with bold lines. The color of nodes represents the clustering label. Most of the nodes in the subgraph belong to the same cluster as the anchor node, indicating conST can aggregate information from spots with similar attributes. **d**. GNNExplainer can further support the identified active L-R pairs in cell-to-cell interaction. All the spots are selected from the same cluster c10 of BRCA in **b**. We set a threshold of interaction value to define spots as active in yellow, and inactive in green. The nodes in the identified subgraph of node 3278 all have active L-R pairs interaction as node 3278.

We demonstrate how to analyze the tumour microenvironment for evaluating neighbor-spreading risk with the above-mentioned downstream tasks, which is divided into cross-clusters CCI analysis and within-cluster CCI analysis.

#### 3.6.1 Cross-clusters: Morphotype Prediction

By inputting the cluster prediction, trajectory and pseudotime inferred from our embeddings to TraSig (43), an analyzing tool for cell-to-cell interaction, we identify the interacting cell-type pairs and active ligand-receptor (L-R) pairs for the tumour tissue.

Cluster c2 and c10 demonstrate more active breast-cancer-related genes interactions, such as BRCA1, BRCA2, BRCC3, and TP53 (50). Cluster 6, as an inactive region, represents the tumour edge. Also, according to GO enrichment analysis (Figure 5a) obtained from TraSig, c10 performs the highest potential on cell migration, cell motility, and regulation of the multicellular organismal process. Strong locomotion ability is one of the most representative phenomena for the IDC region to be distinguished from DCIS regions (48). Therefore, we deduce that clusters 2, 6, and 10 are IDC region, tumor edge, and DCIS region, respectively.

#### 3.6.2 Within-cluster: Neighbor-spreading Risk Evaluation

With the property of breast cancer invasive progression, the within-cluster analysis focuses on the interaction-specific function on cell migration, locomotion, and cell proliferation as shown in the GO term analysis (Figure 5a). We used label transfer from Seurat (13) to annotate spots of the given tissue and obtain the probability distribution of spots related to inferred clusters. Then, we scored the spatial L-R co-expression at every spot and its neighbors.

##### IDC Region (c2)

For the IDC region, cell migration and invasion related to miR-205-3p have been recently reported (51). Upregulated LINC00645 significantly influences the progression of cells *in vivo* as miR-205-3p was a target of LINC00645 and LINC00645, modulating TGF-*β*-induced cells locomotion via miR-205-3p (52). Here we detect the interaction between SMAD5, which belongs to the TGF-*β* superfamily of modulators, and LINC00645. As Figure 5b (iv) shows, the interaction is much more active in IDC regions.

Also, it is reported that at least 50% of breast tumours have an activated type 1 insulin-like growth factor-1 receptor (IGF-1R). The active interaction of IGF-1R with its two natural ligands, insulin-like growth factor-1 (IGF-1) and IGF-2, has been associated by many investigations, as one primary risk factor in breast cancer (53). The receptor system is complex since IR and IGF-1R genes can form several types of hybrid receptors. Here we detect the activities of some highly expressed IGFs, such as IGFBP2-IGF1, IGFBP2-IGF1R, and IGFBP2-TUBGCP5 (Supplementary Figure S11). It also performs not only an upregulated expression within IDC spots but also a more active interaction than DCIS and tumor edge regions.

With a higher chance, LINC00645 and IGF families are potentially well-applied targets in clinical trials on breast cancer, especially for IDC region.

##### Tumor Edge Region (c6)

Genes are considered as “amplified” when the ratio of their copy number in tumour and normal samples is greater than two. Estrogen receptor gene amplification is very frequent in breast cancer. It is reported that more than 20% of breast cancers harbor genomic amplification of the ESR1 gene1 (54). We detect the interaction of some top amplified genes such as ERBB2 (Figure 5b (iii)), ESR1, and CCND1 (Supplementary Figure S12). The result shows that these genes, such as ERBB2-CCND1, have a very active interaction on the border of tumor edge (c2) and IDC region (c6). The overexpression of these amplified gene enhances metastasis-related properties (invasion, angiogenesis, increased survival) of cancer cells that might lead to increased cancer metastases (55). It is very possible that cancer cells from the IDC region are invading the tumor edge, and the tumor edge area will decrease after several cell cycles.

##### DCIS Region (c10)

We further evaluate the following stage of the DCIS region. RAB4A is an essential regulator as it is functional for the fast recycling of integrin *β*3. Integrin *β*3 regulates cell polarity and migration when localized appropriately to the plasma membrane, thereby having an essential role in cancer metastasis (56). Also, GGA2 functions in recycling endosomes to retrieve endocytosed EGFR, thereby sustaining its expression on cell surface, and consequently, cancer cell growth (48). We detect the interaction of LR-pairs RABEP1-GGA2 (Figure 5b (ii)), RABEP1-GGA1, RABEP1-RAB4A, RABEP1-RAB5A, *etc* (Supplementary Figure S13). These pairs show a more obvious interaction, with a strong possibility indicating that the DCIS region is deteriorating and heading to the next stage: Invasive Ductal Carcinoma.

Knowing when a lesion will or will not be life-threatening is essential and requires a thorough understanding of the tumour microenvironment and cancer progression. Here, with the learned embeddings from conST, we can perform more accurate evaluations of the cell-to-cell interactions about IGF-related genes, top amplified genes, and cell metastasis. We conclude the high risk on neighbor-spreading for the IDC region in this special case and ensure that the outcome knowledge will contribute to the decision-making for clinicians in general.

### 3.7 Interpretability

#### 3.7.1 Clustering Explanation

While conST demonstrates strong performance on clustering tasks, we are also interested in why it clusters a specific spot into a specific layer, *i*.*e*., which neighboring spots contribute most to the prediction. Thus, given a spot, we utilize GNNExplainer to identify a subgraph that contributes most to the prediction results.

In Figure 5c, we select nodes 971 and 1765 near the cluster boundary of slice 151673 in spatialLIBD for illustration, where the subgraphs are plotted in bold lines and node color represents the predicted clustering labels. In Figure 5c (i), all the nodes in the subgraph are in the same cluster with node 971, suggesting node 971 is predicted based on the node attributes of those nodes. In Figure 5c (ii), although most of the nodes in the subgraph are in the same cluster with node 1765, node 1577 is in the other cluster. However, the ground truth label shows node 1577 and node 1765 are in the same layer, which means the subgraph we generate is reasonable. Thus, when the subgraph does not match the predictions, it provides further information and helps to identify the reason, which would be helpful in practice. Also, it demonstrates that conST is still able to learn how to aggregate the information and recognize the underlying important relationship among spots in difficult decision boundaries. The use of GNNExplainer points out a way to better refine the network.

#### 3.7.2 CCI Explanation

GNNExplainer can also provide interpretability to CCI on a more fine-grained scale. We select spots all from the same cluster c10 of BRCA (Figure 5b (i)) and inspect the interaction in IDC of pair SMAD5-LINC00645 (Figure 5b (iv)). We set a threshold of interaction value to define spots as active in yellow, and inactive in green (Figure 5d). Setting node 3278 as input, the nodes involved in the subgraph all have active L-R interactions. It demonstrates that conST can not only identify clusters, but also the predictions made by conST are supported by the realistic biological process.

## 4 Conclusion

We propose conST, a powerful and flexible framework for SRT data analysis with contrastive learning. conST can learn low-dimensional embeddings by effectively exploring the multiple modalities of SRT data, including gene expression, spatial information, and morphology (if accessible). The learned embeddings can be utilized in various downstream tasks and the performance surpasses other concurrent methods. Furthermore, we utilize conST to study the tumour microenvironment and cell-to-cell interaction of a breast cancer dataset in detail. conST reveals the cancer development stages and ligand-receptor pairs that reflect the cancer progression. It proves that conST can be used in more complex scenarios, providing more insights for future treatment.

The GNNExplainer explains which neighboring spots contribute to the prediction that conST makes, which is also biologically consistent with the interaction of the L-R pair identified in CCI. The interpretability will enable conST to be used in complex clinical situations with more convincing predictions.

## 5 Acknowledgements

This work was supported by the United Kingdom Research and Innovation (grant EP/S02431X/1), UKRI Centre for Doctoral Training in Biomedical AI at the University of Edinburgh, School of Informatics.

