## Supplementary Material for "conST: an interpretable multi-modal contrastive learning framework for spatial transcriptomics"

This supplementary material provides detailed descriptions of some of the methods mentioned in the paper, the implementation for reproducibility, and more experimental results.

### 1. DETAILED EXPLANATION OF MAE

MAE [1] is a self-supervised computer vision algorithm for representation learning. The framework of MAE is shown in Figure S1. MAE divides the input image to small non-overlapping patches, and then randomly masks a large subset of patches. The remaining patches are input into the encoder. The encoded patches along with mask tokens are processed by a lightweight decoder to reconstruct the input, where the mask tokens indicates the presence of the masked patches. The encoder part is a vision transformer [2] which achieves remarkable results in feature extraction compared to traditional convolutional neural networks (CNNs). The objective of MAE is to learn the representation by reconstructing the heavily masked input with Mean Square Error (MSE).

The use of MAE to extract features from histology images has following advantages over using pretrained ResNet: 1) In terms of statistics, MAE achieves better performance on nearly all commonly used computer vision benchmark datasets than ResNet, demonstrating a much stronger ability for feature extraction. 2) Self-supervised model like MAE learns representation by designing proxy tasks without manual labels. Therefore, the feature extractor can obtain richer information from the data itself instead of the class label containing less information. Hence, the features learned by MAE are more robust and transferable than ResNet even if they are all pretrained on ImageNet. Using MAE for histology images of SRT data, the extracted features can be consistent with the real morphology and gene expression.

| Dataset | Discription | Statistics | Access |
|---|---|---|---|
| SpatialLIBD [3] (10X Visium) | The data is from portion of the DLPFC. It spans six neuronal layers and white matter. | 12 slices Slice 151673: # spots: 3639 # genes: 33538 | http://spatial.libd.org /spatialLIBD/ |
| BRCA [4] (10X Visium) | Fresh frozen Invasive Ductal Carcinoma breast tissue. | # spots: 3798 # genes: 36601 | https://www.10xgenomics.com/ resources/datasets/human-breast-cancer- block-a-section-1-1-standard-1-1-0 |
| MERFISH [5] | Part of the preoptic region of hypothalamus in mouse brain | # spots: 6412 # genes: 155 | https://github.com/haotianteng /FICT-SAMPLE |
| seqFISH [6] | mRNAs in single cell with high accuracy and sub-diffraction-limit resolution-in the cortex | # spots: 913 # genes: 10000 | https://www.spatialomics.org/SpatialDB /seqfish_30911168_browse.php |
| Stereo-seq [7] | Data from mouse olfactory bulb tissues | # spots: 19527 # genes: 27106 | https://github.com/JinmiaoChenLab /SEDR_analyses |

**Table S1.** Detailed description of the datasets experimented in the paper.

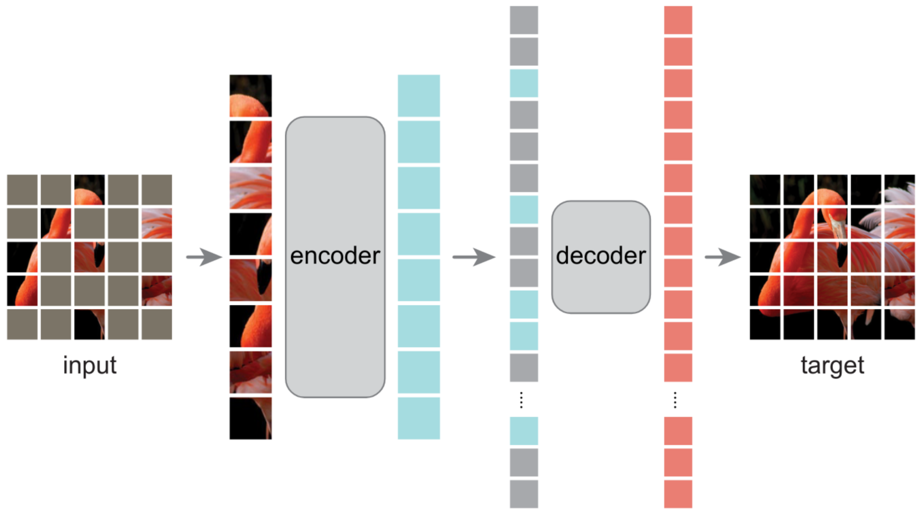


**Fig. S1.** Framework of MAE. MAE randomly masks a large subset of patches, which are divided from the input image. Then, the remaining patches are input into the encoder and reconstructed by decoder where mask token is utilized. MAE learns the robust representation by reconstructing the heavily masked input. Figure reused from [1].

### 2. EXPERIMENTS

#### A. Implementation details

In this section, we provide the implementation details for easier reproducibility and fair comparison. The description and statistics of datasets we used to verify conST is provided in Table S1. Also, we provide the version and access link to the software package in Table S2.

The original SpaGCN does not have the function of generating embeddings, we modified their source code of *predict* function from line 66 of *Spagcn.py* to return the embedding values after the first graph convolutional layer for fair comparison.

#### B. Metrics

First, we introduce several metrics for evaluation of the clustering results. ARI is used for data with tissue/cell type labels, e.g. spatialLIBD. Silhouette Coefficient (SC), Calinski Harabasz Score (CHS), and Davies Bouldin Index (DBI) is

| Method | Version | Language | Access | Ref |
|---|---|---|---|---|
| Monocle3 | 1.0.0 | R | https://cole-trapnell-lab.github.io/monocle3/ | [8] |
| scanpy | 1.8.1 | Python | https://scanpy.readthedocs.io/en/stable | [9] |
| scvelo | 0.2.4 | Python | https://scvelo.readthedocs.io/ | [10] |
| velocyto | 0.17.17 | Python | http://velocyto.org/ | [11] |
| Harmony | 0.1.0 | R | https://portals.broadinstitute.org/harmony/ | [12] |
| Trasig | 1.0.0 | Python | https://github.com/doraadong/TraSig | [13] |
| SpaGCN | 1.2.0 | Python | https://github.com/jianhuupenn/SpaGCN | [14] |
| SEDR | - | Python | https://github.com/JinmiaoChenLab/SEDR | [15] |
| stLearn | 0.3.2 | Python | https://stlearn.readthedocs.io/en/latest/ | [16] |
| BayesSpaces | 1.5.1 | R | https://github.com/edward130603/BayesSpace | [17] |
| Giotto | 1.1.0 | R | https://rubd.github.io/Giotto_site/ | [18] |
| Seurat | 3.2 | R | https://satijalab.org/seurat/ | [19] |

**Table S2.** Detailed descriptions of the software packages used in the paper.

used for data without tissue/cell type labels.

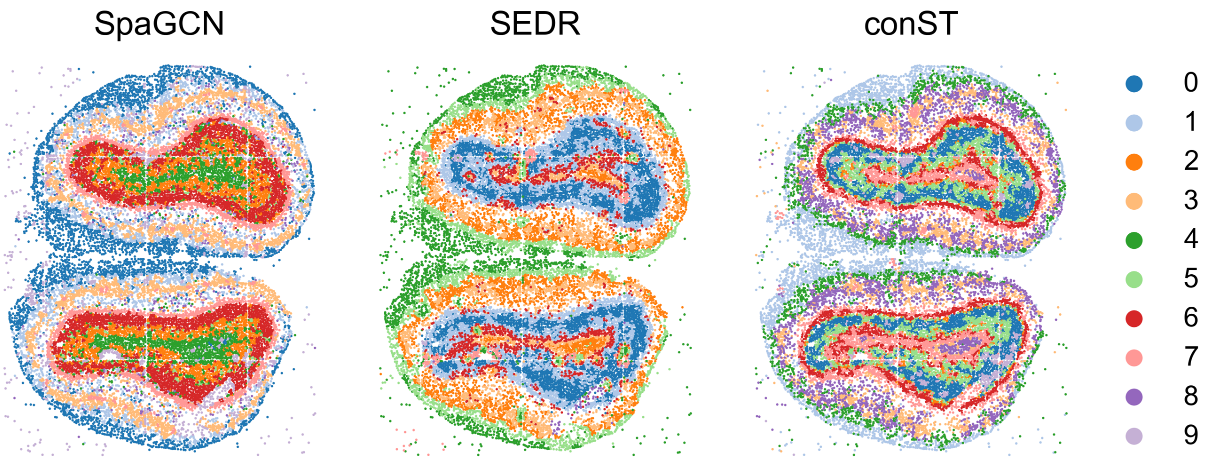


**Fig. S2.** Comparison on Stereo-seq dataset between SpaGCN, SEDR and conST. The boundaries between different clusters, i.e. the inner and outer rings, produced by conST are much clearer than other two methods.

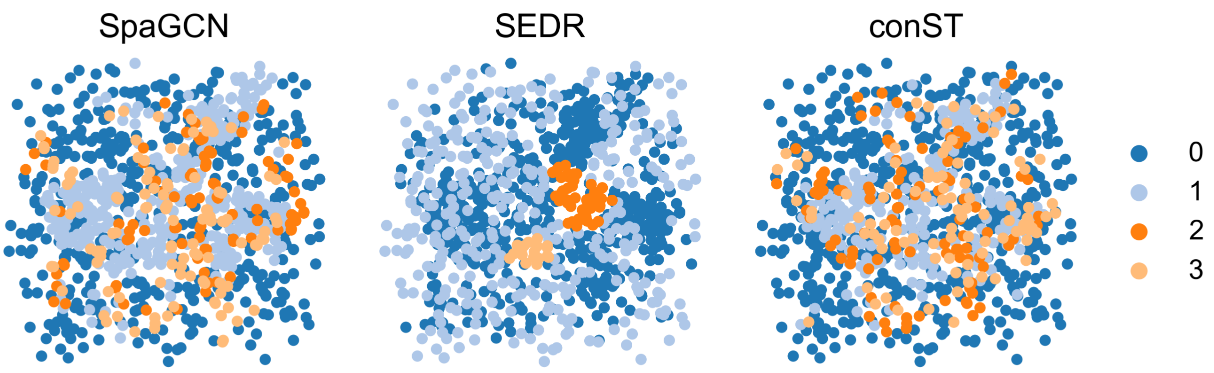


**Fig. S3.** Comparison on seqFISH dataset between SpaGCN, SEDR and conST.

**ARI** Adjusted rand index (ARI) is used to measure the similarity between the predicted labels and ground truth. The Rand Index (RI) calculates a similarity measure between two clusterings, taking all pairs of samples into consideration. It counts pairs that are assigned in the same or different clusters in the predicted and actual clusterings. ARI is a corrected-for-chance version of the Rand index defined as

$$\mathrm{ARI} = \frac{\mathrm{RI} - \mathbb{E}[\mathrm{RI}]}{\max[\mathrm{RI}] - \mathbb{E}[\mathrm{RI}]}, \text{ with } \mathrm{RI} = \frac{\mathrm{TP} + \mathrm{TN}}{\mathrm{TP} + \mathrm{TN} + +\mathrm{FP} + +\mathrm{FN}} \tag{S1}$$

where $\mathbb{E}$ is the expectation value.

**SC** Silhouettes Coefficient (SC) is a metric used to evaluate the clustering effectiveness on dataset without ground truth label [20]. $a_i$ is denoted as tightness, which represents the average distance between sample $i$ and other samples within same cluster. The average distance between a sample and other samples from different cluster is defined as separation. $b_i = \min[b_{i1}, b_{i2}, ..., b_{ik}]$, which is the separation of sample $i$. Then the Silhouette Coefficient (SC) of sample $i$ is defined as

$$s(i) = \frac{b(i) - a(i)}{\max\{a(i), b(i)\}} \tag{S2}$$

The SC score is the average of $s(i)$ ranging from $-1$ to 1 and higher score means higher performance.

**CHS** Calinski-Harabasz score (CHS) [21] is used to evaluate the model without ground truth label. We denote $C_q$ as $q$th cluster, $k$ as the number of clusters, $n_q$ and $c_q$ are the number of points and centroid of the $q$th cluster respectively. $c$ is the global centroid and N is the number of cluster. It is colculated as

$$s(k) = \frac{Tr(B_k)}{Tr(W_k)} \times \frac{N-k}{k-1}, \text{ with } W_k = \sum_{q=1}^{k} \sum_{x \in C_q} (x - c_q)(x - c_q)^T, \quad B_k = \sum_q n_q (c_q - c)(c_q - c)^T \tag{S3}$$

$B_k$ and $W_k$ are representing between-clusters dispersion mean and within-cluster dispersion respectively. Higher CH score means better clustering performance.

**DBI** The Davies-Bouldin index (DBI) [22], which is an internal evaluation scheme, can be used to evaluate clustering algorithm. Assuming we have n cell spots, let $C_j$ be the $j$th cluster of spots. We have $X_j$ as an $n-$dimensional feature vector representing a vectoer of a cell spot assigned to cluster $C_j$.

$$S_i = \left( \frac{1}{T_i} \sum_{j=1}^{T_i} \left| \left| X_j - A_i \right| \right|^p \right)^{1/p} \tag{S4}$$

Here $A_i$ is the centroid of $C_j$. $T_i$ is the number of spots in cluster $i$. We usually have p equal to 2 to make the distance a Euclidean distance function. $S_i$ represents the separation of cluster $i$.

$$M_{ij} = \left( \sum_{k=1}^{N} \left| a_{ki} - a_{kj} \right|^q \right)^{1/q} \tag{S5}$$

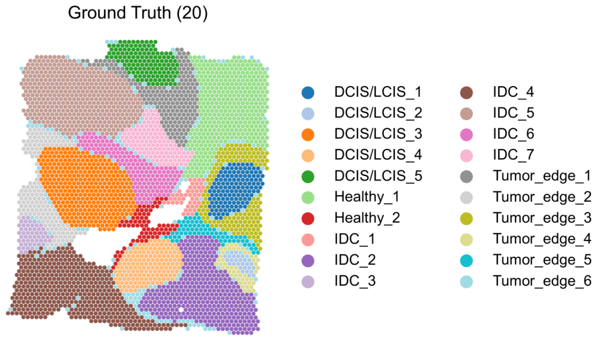


**(a)** Ground-truth label with 20 clusters of BCRA dataset in terms of histology features.

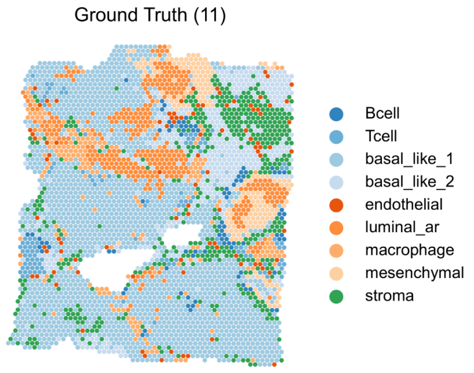


**(b)** Ground-truth label with 11 clusters of BCRA dataset in terms of immune cell types.

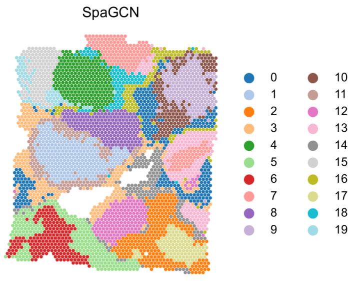


**(c)** Inferred clustering result from the embeddings of SpaGCN.

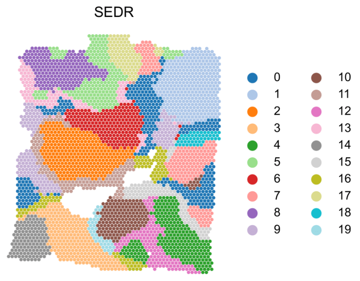


**(d)** Inferred clustering result from the embeddings of SEDR.

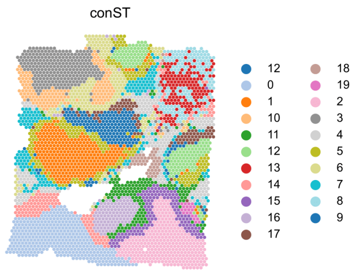


**(e)** Inferred clustering result from the embeddings of conST.

**Fig. S4.** We compare two ground-truth annotations with the results of SpaGCN, SEDR, and conST of BRCA dataset. The 20-cluster ground truth (a) are labeled according to histology features, while the 11-cluster ground truth (b) are labeled according to the immune cell type. Visually, the clusters predicted by conST best aligns both two ground truth labels. conST successfully clusters intact cancer and tissue types, while the clusters of SpaGCN and SEDR are not mixed. Also on the top right corner where there are rich B cell and T cell, the cluster pattern (cluster 8 and 13 in (e)) is also coherent with the immune activities.

Where $a_{ki}$ is the $k$th element of $A_i$, which is the centroid of cluster $i$. Then we have DBI score

$$DBI = \frac{1}{N}\sum_{i=1}^{N} \max_{j \neq i} \left( \frac{S_i + S_j}{M_{ij}} \right) \tag{S6}$$

The DBI score is greater than or equal to zero and lower score means better performance.

### C. Results

Here we list concrete results for various datasets:

For spatialLIBD dataset [3], we provide the clustering ARI of each slice in Table S3.

For Human Breast Cancer (Block A Section I), in short as BRCA, we compare two ground-truth annotations from [15] and [16] with the results of SpaGCN, SEDR, and conST (Figure S4). The 20-cluster ground truth (Figure S4a) are labeled according to histology features, while the 11-cluster ground truth (Figure S4b) are labeled according to the immune cell type. Visually, the clusters predicted by conST best aligns both two ground truth labels. conST successfully clusters intact cancer and tissue types, while the clusters of SpaGCN and SEDR are not mixed. Also on the top right corner where there are rich B cell and T cell, the cluster pattern (cluster 8 and 13 in Figure S4e) of conST is also coherent with the immune activities.

| Slice ID | # Clusters | conST | SEDR | SpaGCN | stLearn | BayesSpace | Giotto | Seurat |
|---|---|---|---|---|---|---|---|---|
| 151507 | 7 | 0.460 | 0.425 | 0.425 | 0.335 | **0.467** | 0.273 | 0.333 |
| 151508 | 7 | 0.337 | 0.308 | 0.362 | 0.307 | **0.437** | 0.213 | 0.380 |
| 151509 | 7 | **0.488** | 0.379 | 0.460 | 0.432 | 0.382 | 0.302 | 0.250 |
| 151510 | 7 | 0.388 | 0.403 | **0.447** | 0.445 | 0.434 | 0.31 | 0.286 |
| 151669 | 5 | 0.458 | 0.303 | 0.226 | 0.328 | **0.469** | 0.321 | 0.328 |
| 151670 | 5 | **0.446** | 0.358 | 0.372 | 0.208 | 0.430 | 0.347 | 0.214 |
| 151671 | 5 | 0.491 | 0.487 | 0.555 | 0.389 | **0.733** | 0.391 | 0.221 |
| 151672 | 5 | **0.62** | 0.494 | 0.439 | 0.56 | 0.347 | 0.253 | 0.187 |
| 151673 | 7 | **0.65** | 0.557 | 0.516 | 0.308 | 0.550 | 0.361 | 0.375 |
| 151674 | 7 | **0.546** | 0.436 | 0.323 | 0.411 | 0.299 | 0.379 | 0.404 |
| 151675 | 7 | **0.507** | 0.464 | 0.300 | 0.393 | 0.53 | 0.260 | 0.292 |
| 151676 | 7 | **0.447** | 0.393 | 0.345 | 0.373 | 0.368 | 0.267 | 0.311 |
| Median | | **0.474** | 0.414 | 0.399 | 0.381 | 0.436 | 0.306 | 0.3015 |
| Mean | | **0.487** | 0.417 | 0.398 | 0.374 | 0.454 | 0.306 | 0.298 |

**Table S3.** Concrete ARI of each slice in spatialLIBD dataset. Numbers in bold indicates the best performance among the compared methods.

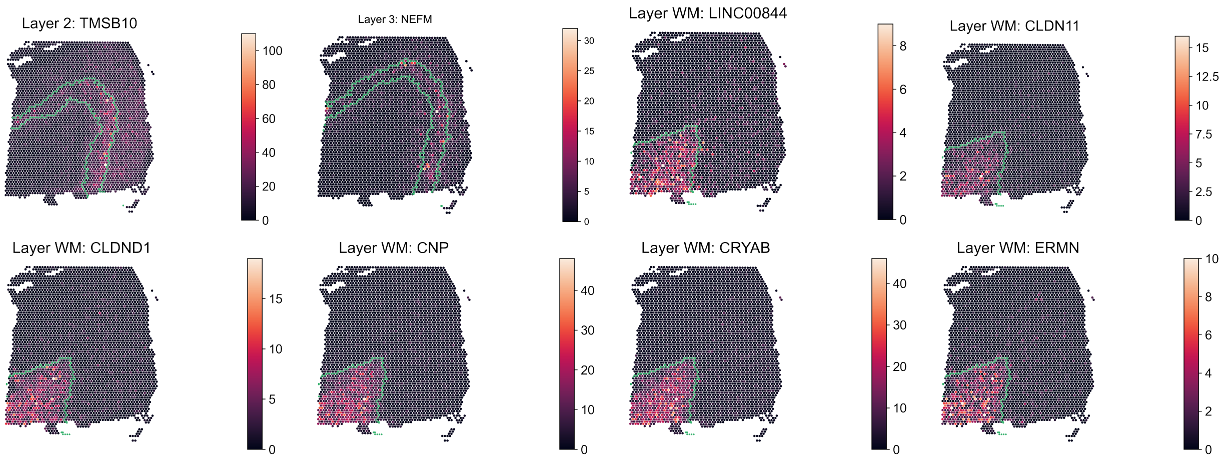


**Fig. S5.** Examples of SVGs detected by conST in spatialLIBD dataset. It can be seen that the highly expressed area aligns well with the clustering boundaries in green lines.

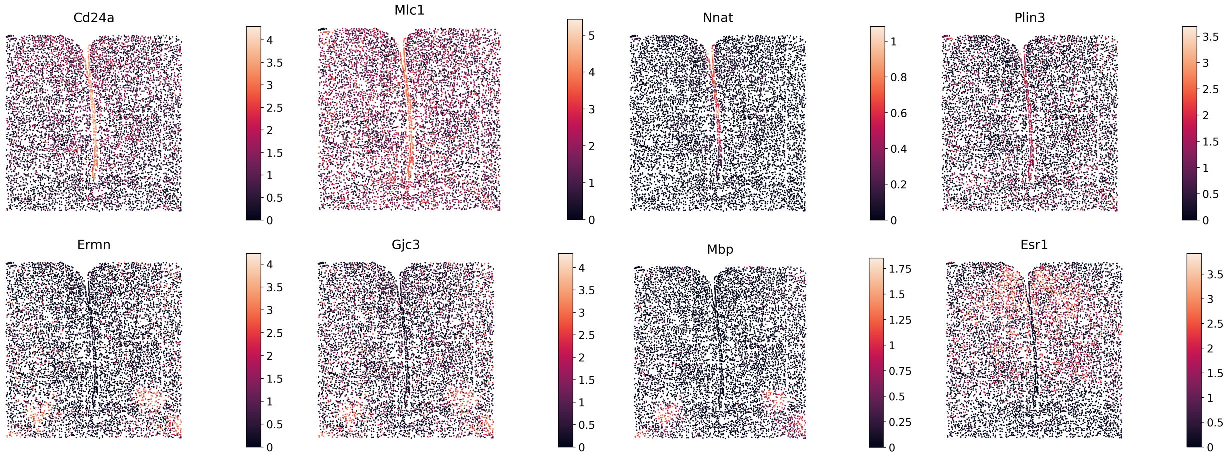


**Fig. S6.** Examples of SVGs detected by conST in MERFISH dataset. The highly expressed areas fit the clusters produced by our algorithm. We can see that *Cd24a*, *Mic1*, *Nnat*, *Plin3* are enriched in the middle line domain. *Ermn*, *Gjc3*, *Mbp* are highly expressed in bottom-left and bottom-right parts.

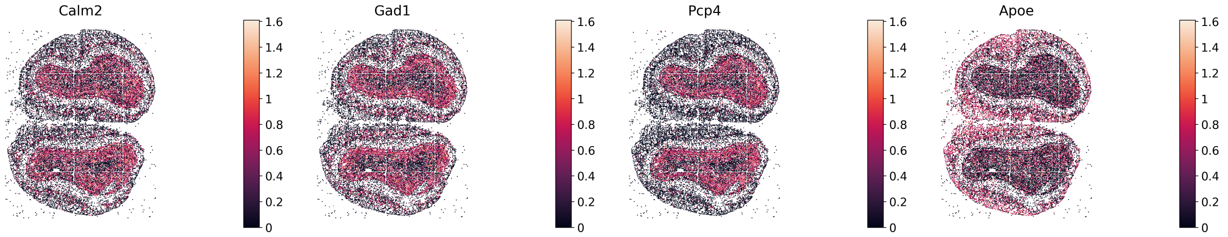


**Fig. S7.** Examples of SVGs detected by conST in Stereo-seq dataset. We can see that the detected genes are highly expressed in the internal ring and external ring, which aligns well with the clusters. Especially, the expression of *Pcp4* is located in the internal ring precisely, which indicates that SVGs detected by conST can well mark the domain where we detect it.

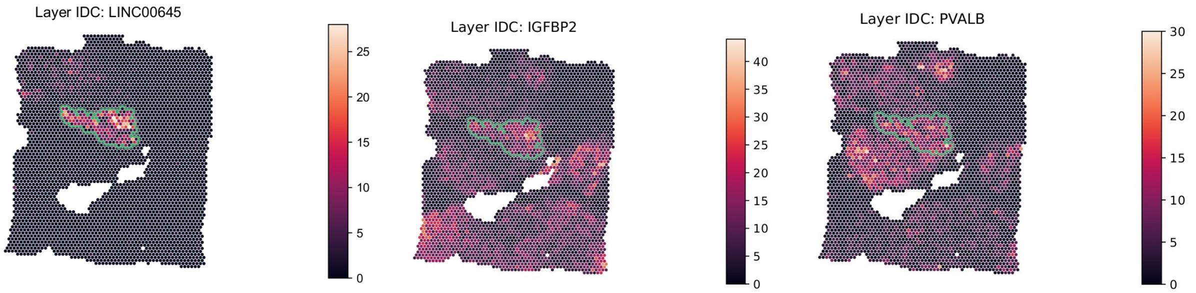


**Fig. S8.** Examples of SVGs detected by conST in the IDC region of BRCA dataset. The highly expressed areas can fit the clusters produced by our algorithm.

| Methods | SC | DBI | CHS |
|---|---|---|---|
| SpaGCN | 0.5 | 2.9 | 260 |
| SEDR | 0.3 | 2.6 | 101 |
| conST | **0.8** | **1.9** | **603** |

**Table S4.** conST outperforms other two methods on merfish datatset evaluated by three metrics. Note that higher SC score and CHS means better effectiveness of clustering and lower DBI means better performance.

| Methods | SC | DBI | CHS |
|---|---|---|---|
| SpaGCN | -0.3 | 26.1 | 18 |
| SEDR | 0.7 | 2.1 | 333 |
| conST | **0.15** | **1.6** | **7956** |

**Table S5.** conST has better performance among these three methods. The SC scores and CHS of conST are much higher than SEDR and SpaGCN, and DBI is much smaller, which indicates that conST is better at clustering task.

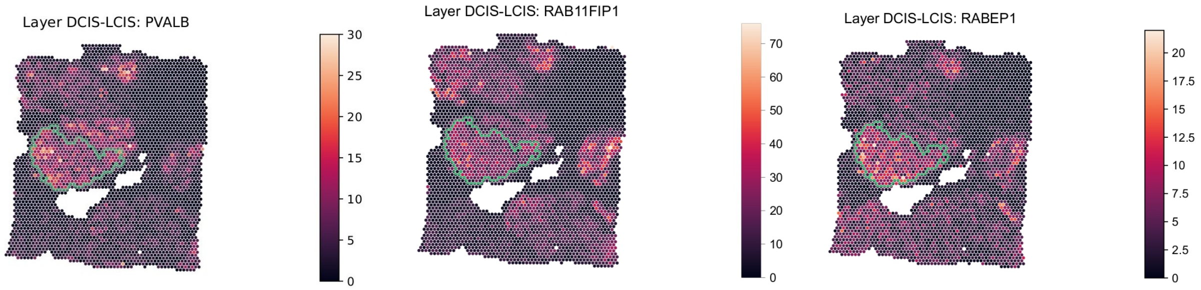


**Fig. S9.** Examples of SVGs detected by conST in the DCIS region of BRCA dataset. It is obvious that the highly expressed areas can basically fit the clusters produced by our algorithm.

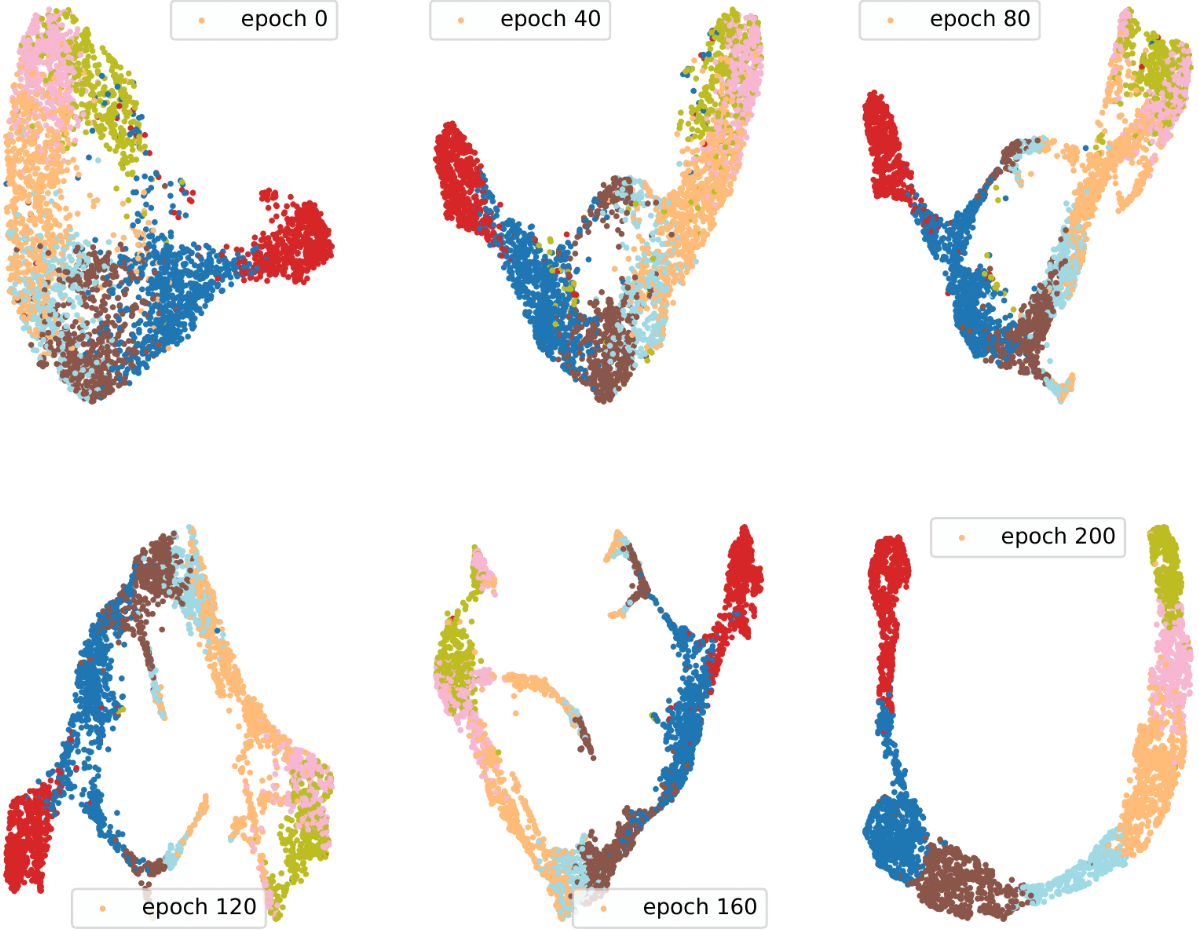




**Fig. S10.** We monitor the training process of slice 151673 in spatialLIBD and visualize the embedding every 40 epochs in during training. It can be seen that the embedding of different clusters are gradually separated through training and converges to a stable state, which demonstrates the effectiveness of conST.

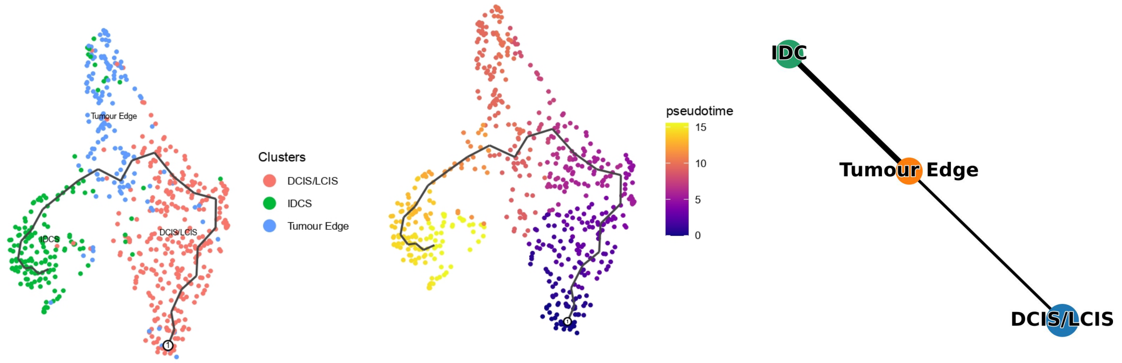


**(a)** Inferred trajectory of BRCA dataset from the embeddings of conST.

**(b)** Inferred pseudotime ordering of BRCA dataset from the embeddings of conST.

**(c)** Inferred trajectory by PAGA of BRCA dataset from the embeddings of conST.

**Fig. S11.** Trajectory inference, pseudotime ordering and PAGA results for BRCA dataset tumor tissue from the embeddings of conST.

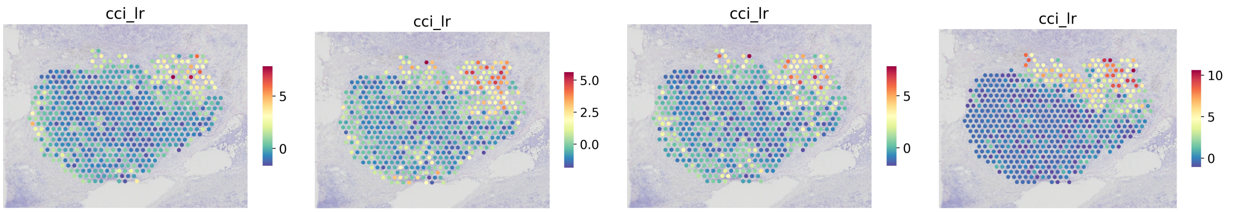


**(a)** IGFBP2-IGF1 **(b)** IGFBP2-IGF1R **(c)** IGFBP2-TUBGCP5 **(d)** SMAD5-LINC00645

**Fig. S12.** Highly expressed interaction pairs in the IDC region.

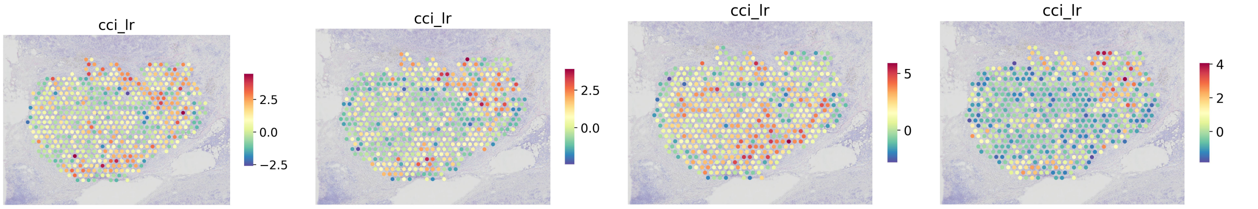


**(a)** ERBB2-CCND1 **(b)** ESR1-CCND1 **(c)** ESR1-ERBB2 **(d)** SPOP-CCND1

**Fig. S13.** Highly expressed interaction pairs in the tumor edge region.

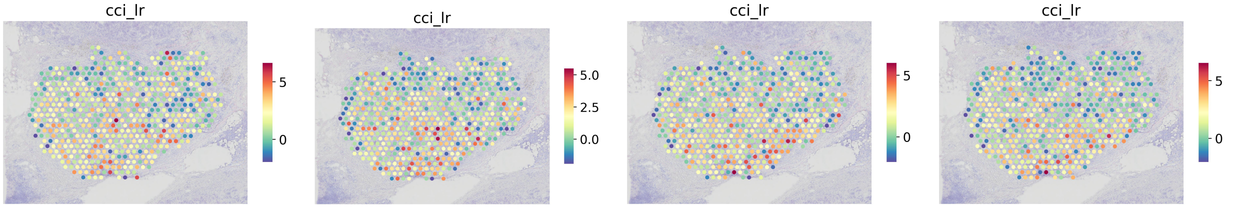


**(a)** RABEP1-GGA1 **(b)** RABEP1-GGA2 **(c)** RABEP1-RAB4A **(d)** RABEP1-RAB5A

**Fig. S14.** Highly expressed interaction pairs in the DCIS region.
